# Ploidy inference from single-cell data: application to human and mouse cell atlases

**DOI:** 10.1101/2023.08.26.554926

**Authors:** Fumihiko Takeuchi, Norihiro Kato

## Abstract

Ploidy is relevant to numerous biological phenomena, including development, metabolism, and tissue regeneration. Single-cell RNA-seq and other omics studies are revolutionizing our understanding of biology, yet they have largely overlooked ploidy. This is likely due to the additional assay step required for ploidy measurement. Here, we developed a statistical method to infer ploidy from single-cell ATAC-seq data. When applied to the data from human and mouse cell atlases, our method enabled systematic detection of polyploidy across a range of cell types. This method allows for the integration of ploidy analysis into single-cell studies.

**Article summary:** Ploidy plays a crucial role in many biological processes. Though modern studies offer deep insights into biology, they often neglect ploidy because it’s challenging to measure. In this research, we have created a new method to detect ploidy using single-cell data. This technique helps identify polyploid cells across various cell types, bridging a gap in our understanding.

## Introduction

While the majority of human and mouse somatic cells are diploid, certain cell types, such as hepatocytes, cardiomyocytes, and megakaryocytes, exhibit higher levels of ploidy (Biesterfeld *et al*. 1994; Ugo 2007; Orr-Weaver 2015; Fang *et al*. 2022). Ploidy is relevant to various biological phenomena, including development, tissue regeneration, metabolism, and cancer. The ploidy level, defined as the number of chromosome sets contained within a cell or nucleus, is termed ‘polyploid’ when it exceeds two. Ploidy can be measured by the DNA content in multiples of the C value, which represents the amount of DNA in a haploid genome. For example, the nuclear DNA content in a diploid or tetraploid cell (in the G1 phase) corresponds to 2C and 4C, respectively.

Historically, cell ploidy has been measured by observing stained DNA or telomeres using fluorescence microscopy or flow cytometry. Although single-cell analysis can incorporate ploidy by adding a flow cytometry step prior to the single-cell assay (Richter *et al*. 2021), such procedures have been relatively uncommon. Here, we propose a statistical algorithm to infer ploidy by analyzing standard single-cell ATAC-seq (scATAC-seq) or single-nucleus ATAC-seq (snATAC-seq) data.

## Results

In an scATAC-seq assay, when focusing on a single site on an autosome, up to two fragments encompassing the site can be observed from a diploid cell, and up to four fragments from a tetraploid cell. The same principle applies to diploid and tetraploid nuclei in the context of a snATAC-seq assay. From a *p*-ploid cell, the number of fragments observable at a single site follows a binomial distribution with *p* trials—this is the foundational observation employed in this study (Figure 1). In practice, we observe only the 5’ ends (on the positive strand of the genome) of ATAC-seq fragments, rather than all sites on the genome, thereby reducing redundant observations.

**Figure 1.**
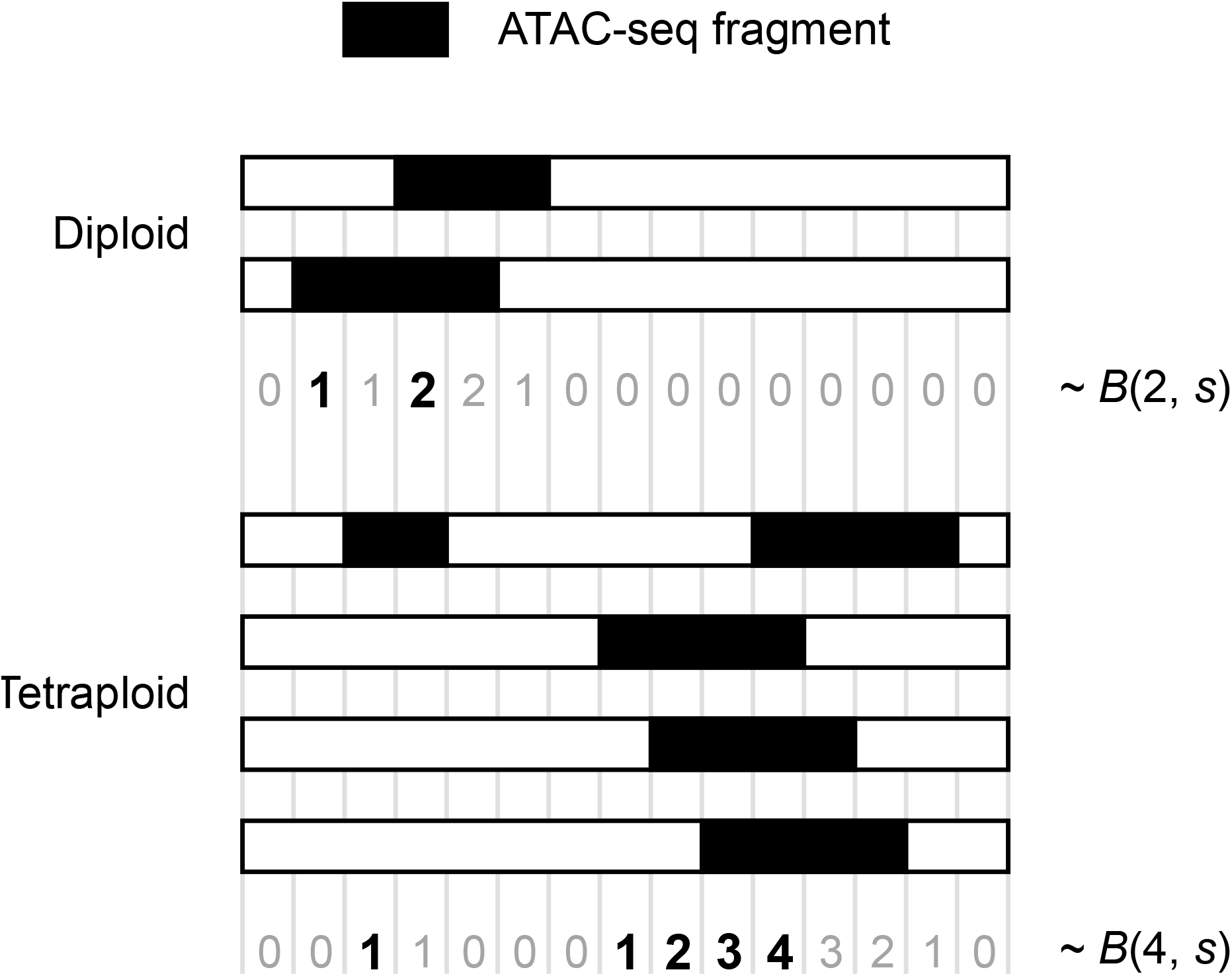
Schematic representation of the probabilistic model employed in this study. At any site on an autosome, the number of snATAC-seq fragments encompassing the site from a diploid cell follows a binomial distribution with two trials and a success probability, *s*. For a tetraploid cell, the number of fragments follows a binomial distribution with four trials. Instead of observing all sites on the genome, we focus only on the 5’ ends (on the positive strand) of the fragments. Observations at the relevant sites are marked in bold black font, while the fragment numbers at ignored sites are indicated in gray font.

We developed two algorithms for ploidy imputation. The first algorithm employs the method of moments for the aforementioned probability distribution and incorporates ploidy as a parameter. In contrast, the second algorithm models fragment counts as originating from distinct distributions corresponding to potential ploidy levels (such as diploid, tetraploid, and octoploid) and infers the cellular affiliation using the Expectation-Maximization (EM) algorithm. If the assumption of the binomial distribution holds true, the first algorithm may perform better. However, if this assumption does not hold, the second algorithm could prove to be superior.

We first evaluated the algorithms using simulated data. We used actual snATAC-seq data for human peripheral blood mononuclear cells (all diploid) and generated tetraploid or octoploid nuclei by combining two or four diploid nuclei, respectively. We generated a mixture of 1521 instances each for diploid, tetraploid and octoploid nuclei, totaling 4563 nuclei. The performance of the algorithms was assessed using 100 randomly generated datasets (Figure 2). Sensitivity, specificity, precision, and negative predictive value for diploids, tetraploids, and octoploids were all notably high when utilizing the method of moments. However, when using the EM algorithm, we found some measurements to be inferior to random guesses. Interestingly, when examining the negative predictive value for diploids, we found that the EM algorithm (mean = 0.982, SD = 0.0025) outperformed the moment method (mean = 0.941, SD = 0.0033). This suggests that when the EM algorithm predicts a nucleus to be polyploid (i.e., not diploid), it tends to be more accurate than the prediction made by the moment method.

**Figure 2.**
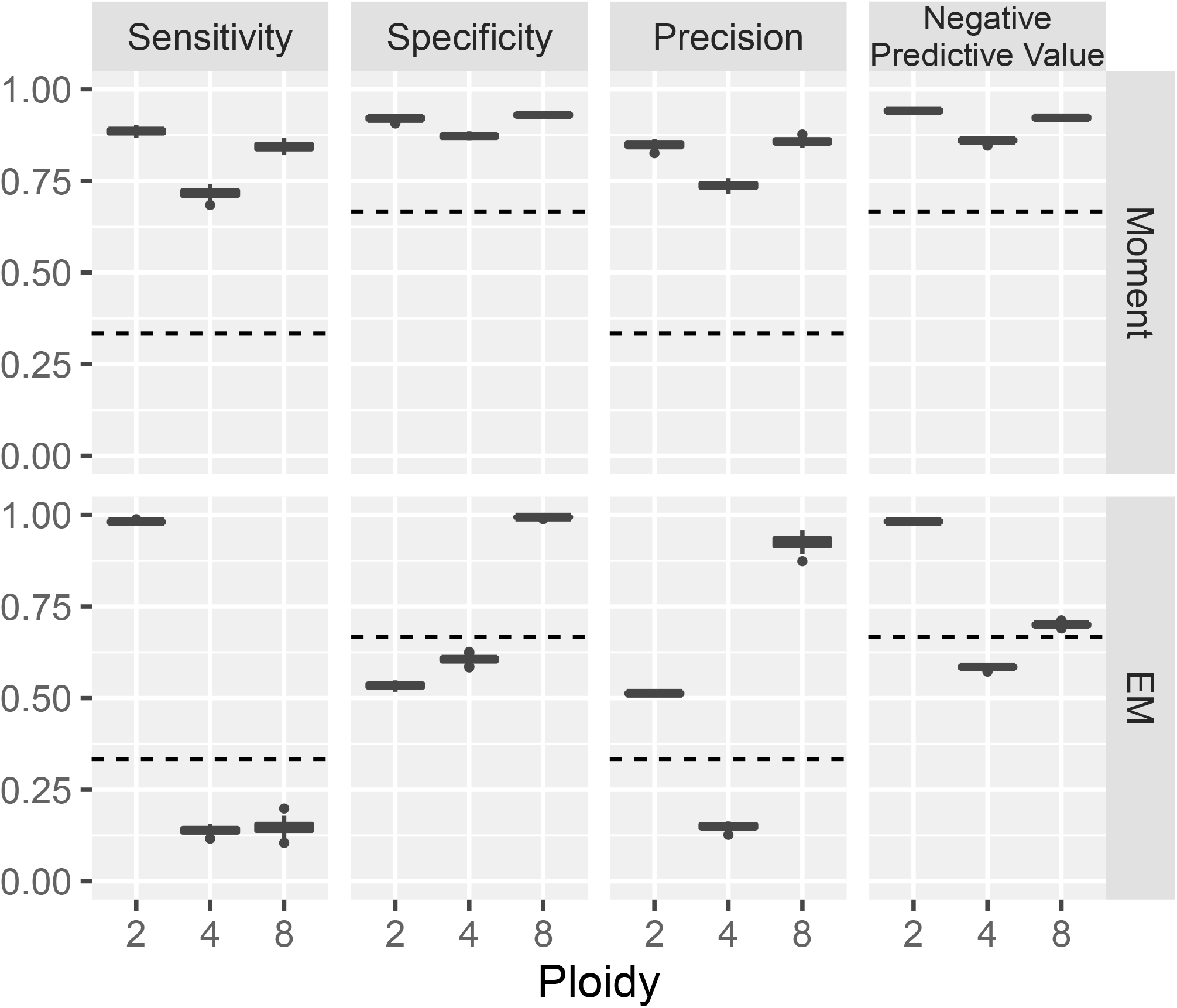
Evaluation of ploidy inference algorithms using simulated data. Sensitivity, specificity, precision, and negative predictive value (vertical axis) for the prediction of diploids, tetraploids, and octoploids (columns) are plotted across 100 datasets simulated from a snATAC-seq experiment on human peripheral blood mononuclear cells. The upper and lower panels represent the method of moments and the EM algorithm, respectively. In the boxplots, the middle bar indicates the median, and the lower and upper hinges correspond to the first and third quartiles, respectively. The whiskers extend to the value no further than 1.5 times the inter-quartile range from the hinges, and data points beyond the hinges are plotted individually. Dashed lines indicate the performance of a random guess.

The performance of the algorithms may decline if the total number of ATAC-seq fragments in an experiment is insufficient. To estimate the effect of next-generation sequencing depth, we conducted simulations with half or a quarter of the reads compared to the original data (57k read-pairs per cell) and observed a limited loss in accuracy (Figure S1). Considering that 25k read-pairs per cell are generally recommended, it is unlikely that the performance of the ploidy inference algorithm will significantly change within the range of usual snATAC-seq experiments.

Next, we applied ploidy inference to publicly available data from cell atlases of the mouse embryo (Jiang *et al*. 2023), human fetus (Domcke *et al*. 2020), adult mouse (Cusanovich *et al*. 2018), and adult human (Zhang *et al*. 2021). To avoid mischaracterization of a diploid nucleus as polyploid, we employed the EM algorithm. We detected polyploids in 22 cell types across 17 tissues (Table 1).

**Table 1.** Polyploid cell types detected in mouse and human cell atlases.

| **Table 1. Polyploid cell types detected in mouse and human cell atlases** |  |  |  |  |  |
| --- | --- | --- | --- | --- | --- |
| Tissue | Cell type | Mouse embryo | Human fetus | Adult mouse | Adult human |
| *Mesoderm derived* |  |  |  |  |  |
| Heart | Cardiomyocyte | n.s. | 2.8E-306 | 1.8E-76 | 8.7E-251 |
| Heart | Mesothelial cell | n.s. | - | - | 5.8E-256 |
| Spleen | B cell | - | - | 2.0E-132 | - |
| Liver | Erythroblast | 0 | n.s. | - | - |
| Kidney | Proximal tubule | - | - | 1.9E-68 | - |
| Gonad female | Premeiotic germ cell | 0 | - | - | - |
| *Endoderm derived* |  |  |  |  |  |
| Liver | Hepatoblast | n.s. | 4.7E-105 | - | - |
| Liver | Hepatocyte | - | - | 0 | n.s. |
| Pancreas | Pancreatic acinar cell | n.s. | - | - | 0 |
| Lung | Endothelial cell | n.s. | 3.8E-281 | n.s. | n.s. |
| Lung | Epithelial cell | 5.6E-96 | 2.4E-95 | - | - |
| Thymus | Thymocyte | - | 0 | - | - |
| Esophagus | Epithelial cell | - | - | - | 2.1E-154 |
| Stomach | Smooth muscle cell | 3.1E-135 | - | - | - |
| Intestine | Epithelial cell / enterocyte | n.s. | 2.8E-130 | n.s. | 1.5E-48 |
| Intestine | Stromal cell | 3.2E-100 | n.s. | - | - |
| *Ectoderm derived* |  |  |  |  |  |
| Brain | Neuron | n.s. | n.s. | 8.7E-216 | - |
| Mammary tissue | Mammary luminal epithelial cell | - | - | - | 2.89E-105 |
| The numbers indicate P-value for the null hypothesis of being diploid. "n.s." indicates not statistically significant. Hyphen "-" indicates not studied in the dataset. |  |  |  |  |  |

The polyploidy of mammalian somatic cells has been well-studied in cardiomyocytes and hepatocytes and has also been reported in other cell types including neurons, mammary gland epithelial cells, and mesothelial cells (Biesterfeld *et al*. 1994; Ugo 2007; Orr-Weaver 2015; Fang *et al*. 2022). We observed polyploidy in these cell types. Moreover, most of the polyploid cell types identified in this study have been previously reported. Polyploidy of tubular cells in the kidney has been reported in both mice and humans (Chiara *et al*. 2022). Polyploidy of pancreatic acinar cells has been reported in mice (Ge and Morgan 1990). Human vascular endothelial cells have been reported to become polyploid during *in vitro* culture (Wagner *et al*. 2001), and pulmonary epithelial cells have been identified as polyploid in both infant and adult humans (Hamada *et al*. 1989). Polyploidy in intestinal epithelial cells has been reported in *Drosophila* (Øvrebø and Edgar 2018). To the best of our knowledge, polyploidy has not been reported in smooth muscle cells in the stomach or in stromal cells in the intestine. Our algorithm could also detect cell types that are actively proliferating, as may be the case for B cells in the spleen and thymocytes in the thymus. The detected polyploids included cell types with biologically apparent roles, such as erythroblasts and premeiotic germ cells in the female gonad.

## Discussion

Currently, the most widely used single-cell assays are RNA-seq, ATAC-seq, and their combined assay. Our ploidy inference algorithm is readily applicable to single-cell ATAC-seq results, with no additional biochemical assays. Through simulations using real data, we demonstrated the high accuracy of our algorithm. When applied to data from cell atlases of humans and mice, we identified a diverse range of polyploid cell types, corroborating previous reports on individual tissues. Although it remains untested, theoretically, our algorithm could also be applied to single-cell whole-genome sequencing. This study is the first attempt to systematically survey polyploidy across a broad spectrum of tissues. Integrating this algorithm into other single-cell studies could shed light on the relationship between ploidy and various biological phenomena at single-cell level.

## Methods

### Probability distribution for the method of moments

In both ATAC-seq and whole-genome sequencing, the nucleotide sequence of chromosomal fragments is read. At any site on an autosome, the number of fragments encompassing that site is at most two from a diploid cell, and at most four from a tetraploid cell. In this context, PCR-duplicates amplified from the same chromosomal fragment are considered identical. Our probabilistic model is formulated based on this straightforward counting of fragments at a site. We assume that the fragments originate from a *p*-ploid cell in the case of single-cell assays, or from a *p*-ploid nucleus in the case of single-nucleus assays.

At one site on an autosome, the number of sequenced fragments should follow a binomial distribution with *p* trials. The probability of observing *x* fragments is

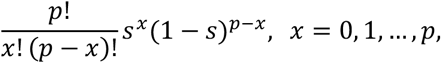

where *s* is the success probability, in this case, the probability of sequencing this site within one chromosome. Observing across all sites on the genome, we count the number of sites where *x* fragments were observed and denote this by *f*_*x*_. We define

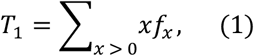

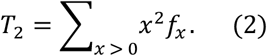

If we could assume that the number of sequenced fragments is independent and identically distributed across the sites, the statistics *T*_1_ and *T*_2_ equal the first and second sample moments, respectively, multiplied by *n*, the number of sites. Hence, the expectations are

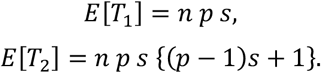

Since

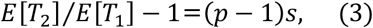

we can use *T*_2_/*T*_1_ − 1 as an estimator of (*p* − 1)*s* as shown by Rider (Rider 1955). The estimator provides information about the ploidy of each cell, multiplied by an unknown value *s* that is constant across cells.

The aforementioned model, however, is practically suboptimal for two reasons: 1) Observations from sites in proximity are statistically dependent, as typically 50–100 bp are sequenced together in a read, and 2) most sites would be covered by zero fragments in the case of ATAC-seq. To address these issues, we alter the observation points from all sites on the genome to the 5’ ends (on the positive strand) of sequenced fragments. Accordingly, we employ a truncated binomial distribution, wherein cases with zero success are ignored. Nonetheless, this modification does not affect the validity of the estimator *T*_2_/*T*_1_ − 1. Firstly, *f*_0_ is not used in the calculation of *T*_1_ and *T*_2_ (equations (1) and (2)). Secondly, although the total number of observation points, *n*, is unknown, it is canceled out and becomes irrelevant in equation (3).

### Inferring ploidy using the method of moments

For the single cells obtained in an experiment, we compute *u* = *T*_2_/*T*_1_ − 1, which is an estimator of (*p* − 1)*s*. In the case of a mixture of diploids, tetraploids, and octoploids, we expect *u* to distribute in three clusters whose values maintain a ratio of (2 − 1): (4 − 1): (8 − 1). We take the logarithm and, for robustness, limit the range: the maximum value is at most twice the 3^rd^ quartile subtracted by the median of the original distribution, while the minimum value is at least twice the 1^st^ quartile subtracted by the median. The obtained value is denoted by *v*_*i*_ for a cell *i*.

Given the set *P* of possible ploidy in a dataset, we first estimate *ŝ* that best fits the data:

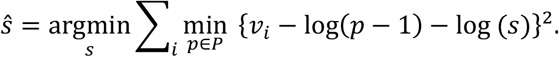

The estimated ploidy of cell *i* is then calculated as

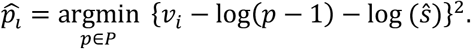

Alternatively, we can estimate ploidy allowing for a fractional value not necessarily in *P* as

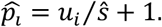

Although we do not explore this further, the fractional ploidy could potentially be used in inferring cell cycle phase.

### Inferring ploidy using EM algorithm

Alternatively, we approach ploidy inference by considering (*f*_1_, *f*_2_, *f*_3_, …) as samples drawn from a mixture of multinomial distributions. The source multinomial distributions correspond to potential ploidy, and the algorithm requires the number of possible ploidies, for instance, 3 for diploids, tetraploids, and octoploids. We impose no restrictions on the multinomial distributions, thus being more permissive than the moment-based method. To assign each cell to a source distribution, we utilize the EM algorithm implemented in the mixtools package (version 1.2.0) (Benaglia *et al*. 2009) of the R software. Based on the analysis of simulated and empirical data, we found the best performance using only the counts (*f*_2_, *f*_3_, *f*_4_), excluding *f*_1_ which is considerably larger than other components and uninformative. In cases where the EM algorithm does not converge, possibly due to *f*_2_ being much larger than the other two components, we use (*f*_3_, *f*_4_, *f*_0_) instead. Ploidy levels (e.g., 2, 4, 8) are assigned to the source distributions according to the size order of the last multinomial parameter, which is for *f*_4_ in the case of (*f*_2_, *f*_3_, *f*_4_).

### Processing of DNA fragment data

Fragments mapped on autosomes were retained unless the 5’-or 3’-end was located within simple repeats of the genome. Fragments in simple repeats were discarded because the mapping of next-generation sequencing reads could be inaccurate. The simple repeats in the mouse and human genomes were obtained from the UCSC table browser (Karolchik *et al*. 2004). The fragment count (*f*_1_, *f*_2_, …, *f*_1_) was computed for each cell.

### Simulated data

To evaluate the performance of the ploidy inference algorithms, we generated a simulated mixture of diploid, tetraploid, and octoploid nuclei. snATAC-seq data for human peripheral blood mononuclear cells from a healthy male was downloaded from the 10x Genomics website (https://www.10xgenomics.com/resources/datasets/10-k-human-pbm-cs-multiome-v-1-0-chromium-controller-1-standard-2-0-0). The dataset included 10,691 nuclei, sequenced to 57k read-pairs per cell. From each cell type, we randomly selected 7 nuclei repetitively. One nucleus was retained as a diploid. ATAC-seq fragments from two of the nuclei were combined and considered as derived from a single tetraploid. The remaining four nuclei were combined and treated as one octoploid. Each simulated experiment consisted of 1521 diploids, tetraploids, and octoploids each, totaling 4563 nuclei. We generated 100 such simulated experiments.

### Cell atlases for mouse and human

We applied ploidy inference to publicly available cell atlases of the mouse embryo (GSA CRA003910), human fetus (GEO GSE149683), adult mouse (GEO GSE111586) and adult human (GEO GSE184462). The datasets comprise 22, 60, 17, and 70 experiments with various tissue samples, respectively. We either downloaded the snATAC-seq fragment data or computed it from FASTQ files using the Cell Ranger ATAC software (version 1.2.0). We retained nuclei with between 10^3^ and 10^5^ fragments. The total number of analyzed nuclei was 202,704 for the mouse embryo, 570,257 for the human fetus, 69,428 for the adult mouse, and 473,467 for the adult human.

For ploidy inference in cell atlases, we employed the EM algorithm for the following reasons. In simulations, the EM algorithm achieved a higher negative predictive value for the diploid, suggesting that the predicted polyploid cells are more likely to be genuinely polyploid. Furthermore, the moment-based method assumes a constant probability of success, *s*, which may not be universally applicable across experiments within a dataset. Ploidy inference was computed for all nuclei in each of the four datasets. Three classes of possible ploidy (diploid, tetraploid and octoploid) were assumed.

Lastly, we sought cell types that had a low proportion of diploids. By plotting the total number of nuclei (horizontal axis) against the proportion of diploids (vertical axis) for cell types in a dataset, we were able to detect highly polyploid cell types as outliers in the bottom right of the plot. Formally, for each cell type in each tissue sample, we empirically tested whether the number of diploid nuclei was smaller than expected. We employed a binomial distribution with the success probability being the average proportion of diploid nuclei in the dataset. We adopted significance levels of 10^−40^ for the adult mouse, 10^−50^ for the human fetus and adult, and 10^−80^ for the mouse embryo, inspecting the outliers in the plots.

## Data availability

The software is implemented as the scPloidy package of the R software and is freely available from CRAN. The source code is available from GitHub (https://github.com/fumi-github/scPloidy). The source code and data for the analysis in this article is available from figshare (https://doi.org/10.6084/m9.figshare.23574066).

## Acknowledgments

This work was supported by the NCGM Intramural Research Fund (20A1013).

