## Supplemental Figure S1 for "Ploidy inference from single-cell data: application to human and mouse cell atlases"

The effect of next-generation sequencing depth on the accuracy of ploidy inference algorithms. In comparison to the simulation in Figure 2, a subsample of 50% or 25% of next-generation sequencing reads (upper panel and lower panel, respectively) was used as input. See the legend of Figure 2 for further details.

Sub-sampling 50% of reads.

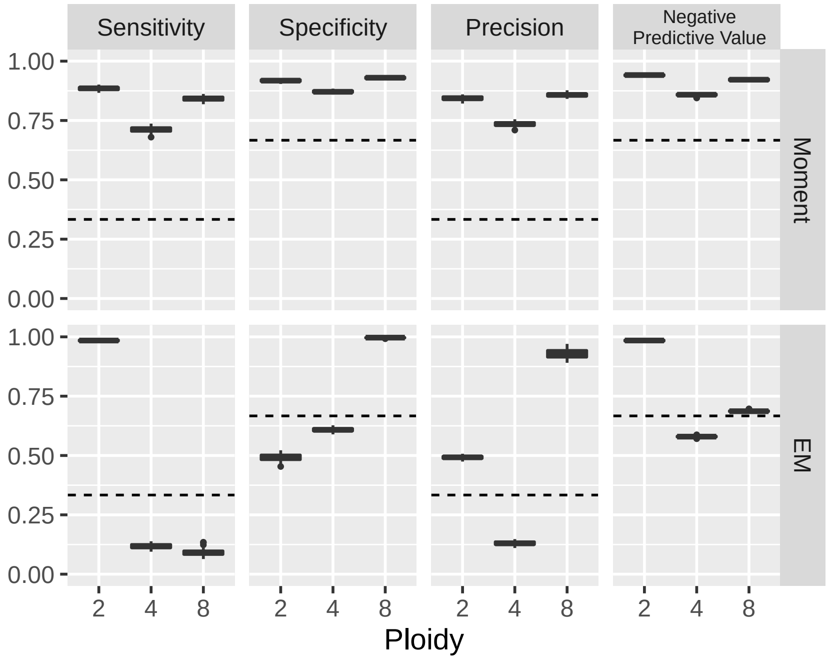


Sub-sampling 25% of reads.

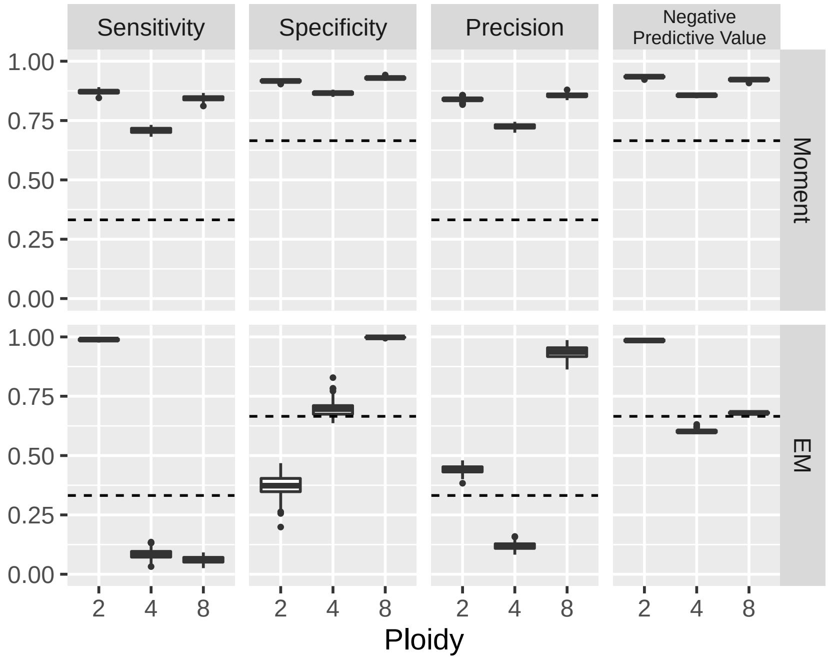
